# *Caenorhabditis elegans* sensory neurons modulate pathogen specific responses via HLH-30/TFEB transcription factor and FSHR-1 GPCR axes of immunity

**DOI:** 10.1101/609610

**Authors:** Anjali Gupta, Manoj Varma, Varsha Singh

## Abstract

Pattern recognition receptors allow animals to sense microbe associated molecular patterns and mount effective immune responses. It is not clear how *Caenorhabditis elegans* recognizes pathogenic microbes in absence of classical pattern recognition pathways. Here, we asked if sensory neurons of *C. elegans* allow it to distinguish between pathogens. Exposure of *C. elegans* to a Gram positive bacterium *Enterococcus faecalis* or to a Gram negative bacterium *Pseudomonas aeruginosa* showed predominantly pathogen-specific signatures. Using nematodes defective in sensory perception, we show that neuronal sensing is essential to mount pathogen specific immune response. OSM-6 expressing, ciliated neurons exert non-cell autonomous control of immune effector production via an OSM-6-FSHR-1 GPCR axis as well as an OSM-6-HLH-30/TFEB transcription factor axis during *E. faecalis* infection. OSM-6-FSHR-1 axis also controls immune response to *P. aeruginosa*. In all, this study delineates essential role of sensory perception in the regulation of pathogen-specific immunity in *C. elegans*.

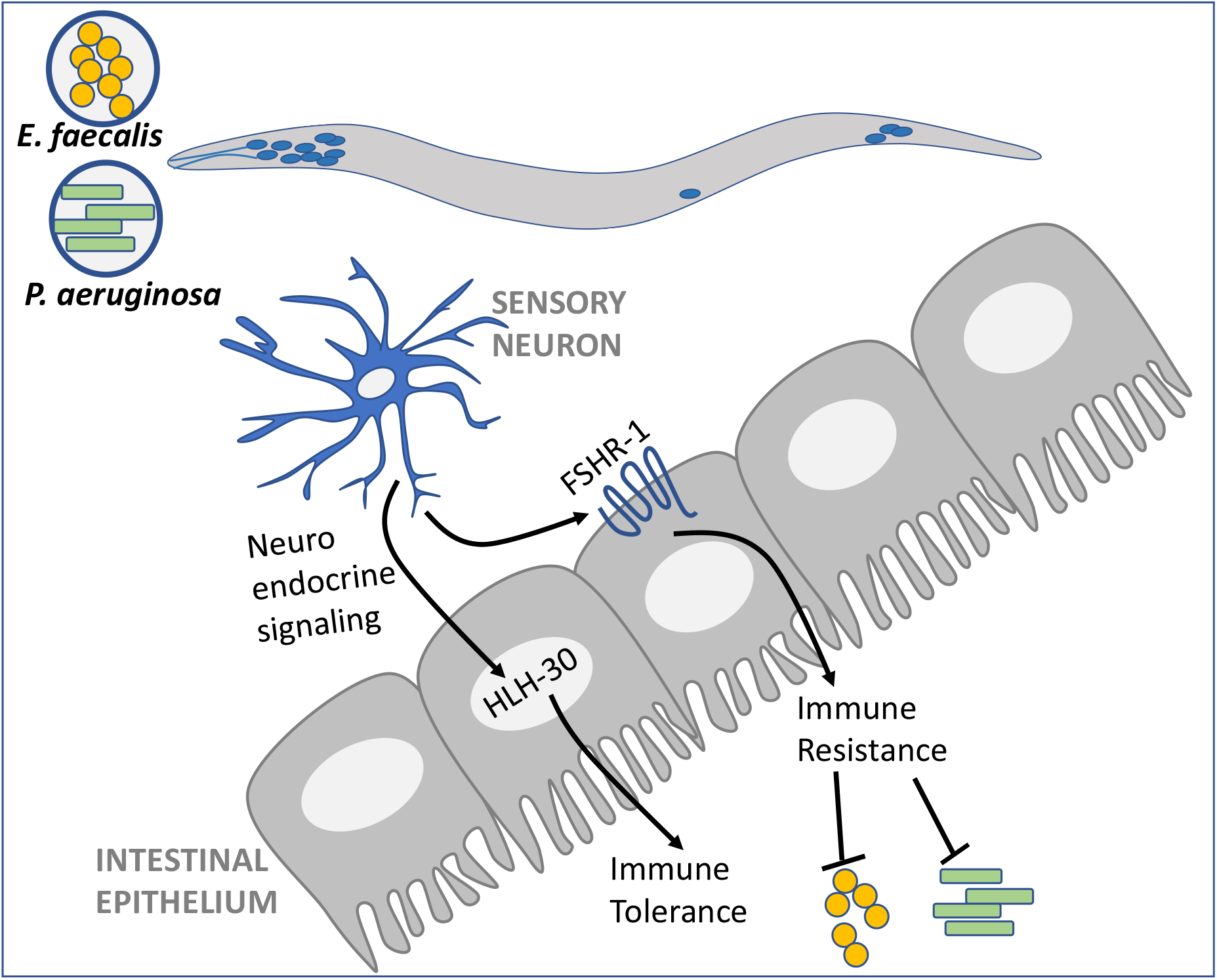

## INTRODUCTION

Appropriate sensing of invading pathogen by the host is central to mounting of an effective immune response and resolution of the infection. Toll and nlr family of pattern recognition receptors (PRRs) have emerged as the central players in pathogen recognition in insects and mammals (Kawai and Akira, 2009). Activation of PRRs results in activation of innate immune responses including antimicrobial peptides (AMPs) and lectin like molecules. *C. elegans* is able to forage for food bacteria in decomposing organic matter containing a number of pathogenic and benign microbes although it lacks components of canonical pattern recognition pathways (Pujol *et al*., 2001; Pradel *et al*., 2007; Tenor and Aballay, 2008). The nematodes can also initiate transcriptional programs for the production of immune effector molecules in response to pathogenic bacteria and fungi (Wong *et al*., 2007; Irazoqui *et al*., 2010; Engelmann *et al*., 2011). Evidence for pathogen specific signalling pathways for survival also exist (Singh and Aballay, 2006; Troemel *et al*., 2006; Estes *et al*., 2010; Richardson, Kooistra and Kim, 2010; Sun, Aballay and Singh, 2016; Marissa Fletcher *et al*., 2019).

There is evidence that G protein coupled receptors might serve as non canonical receptors for sensing pathogenesis. Formyl peptide receptor, FPR-1, in nociceptive neurons directly binds to formylated peptide of *S. aureus* and affects pain response during infection in mice (Chiu *et al*., 2013). In *C. elegans*, NPR-1 and OCTR-1 in the nervous system, FSHR-1 in the intestine, and DCAR-1 in the skin act as regulators of flight response and immune homeostasis on *Pseudomonas aeruginosa* and *D. coniospora* infection (Styer *et al*., 2008; Powell, Kim and Ausubel, 2009; Sun *et al*., 2011; Singh and Aballay, 2012; Zugasti *et al*., 2014; Gupta and Singh, 2017). G protein and beta arrestin signalling in neurons, and endocrine activity by insulin like peptides *ins-7* also regulate survival on *P. aeruginosa* (Kawli and Tan, 2008; Kawli, Wu and Tan, 2010; Singh and Aballay, 2012; Laurenson-Schafer *et al*., 2013; Meisel *et al*., 2014; Lee and Mylonakis, 2017). The nervous system must activate transcription factors at the site of infection, intestine or skin, to facilitate expression of antimicrobial peptides and cytoprotective genes.

Upon infection by a pathogenic microbe, the host can engage in two distinct responses at the site of infection– a) prevent pathogen proliferation (resistance), and (b) protect itself from microbial toxins and host inflammatory response (tolerance) (Tan, 2011). Signalling networks including DAF-2/DAF-16 pathway, p38 MAP Kinase pathway and TGF-β signalling regulate the expression of antimicrobial effectors resulting in clearance of the pathogen and resistance (Troemel *et al*., 2006; Kawli and Tan, 2008; Zugasti and Ewbank, 2009). Neuronal GPCR OCTR-1 and IRE-1/XBP-1 signalling mechanisms regulate immune tolerance during *P. aeruginosa* infection of *C. elegans* (Richardson, Kooistra and Kim, 2010; Sun *et al*., 2011). Previous studies have not examined the contribution of resistance and tolerance mechanisms to survival during infection.

In this study, we set out to ask if sensory neurons of *C. elegans* contribute to pathogen specific immunity to Gram negative and Gram positive bacteria. We show that sensory perception is essential for mounting effective immune response to Gram positive bacterium *Enterococcus faecalis* and to Gram negative bacterium *Pseudomonas aeruginosa*. We show that neuropeptides are essential for immune response to *E. faecalis* but appear dispensable for survival on *P. aeruginosa*. We find that sensory neurons suppress immune response on normal food bacterium *E. coli*, but activate it in the presence of pathogen. Ciliated neurons regulate HLH-30/TFEB transcription factor and FSHR-1 GPCR pathways to regulate both tolerance and resistance mechanism of survival respectively. In all, we show that sensory perception regulates different mechanisms of survival of *C. elegans* in a context/infection specific manner.

## RESULTS

### Exposure to pathogenic bacteria induces pathogen-specific responses in *C. elegans*

Study of early response to pathogens can be instructive in deciphering sensory control of pathogen recognition, immune response and eventual survival. A key question of interest is how *C. elegans* differentiates between different pathogenic bacteria and *E. coli* early on, before pathogen inflicted damage to tissue occur. We analysed nematodes’ early response to infection upon 8 hours exposure to *E. faecalis*, to *P. aeruginosa*, and to *E. coli* OP50. Compared to *E. coli*, a total of 717 transcripts were induced in *E. faecalis* exposed nematodes, while 380 transcripts were induced in *P. aeruginosa* exposed nematodes (Supplementary Table S1, and Table S2). Surprisingly, we found that only 110 genes induced by both the pathogens while majority (607 in *E. faecalis* and 270 in *P. aeruginosa* exposed nematodes) of the genes were pathogen-specific (Fig 1A).

**Figure 1.**
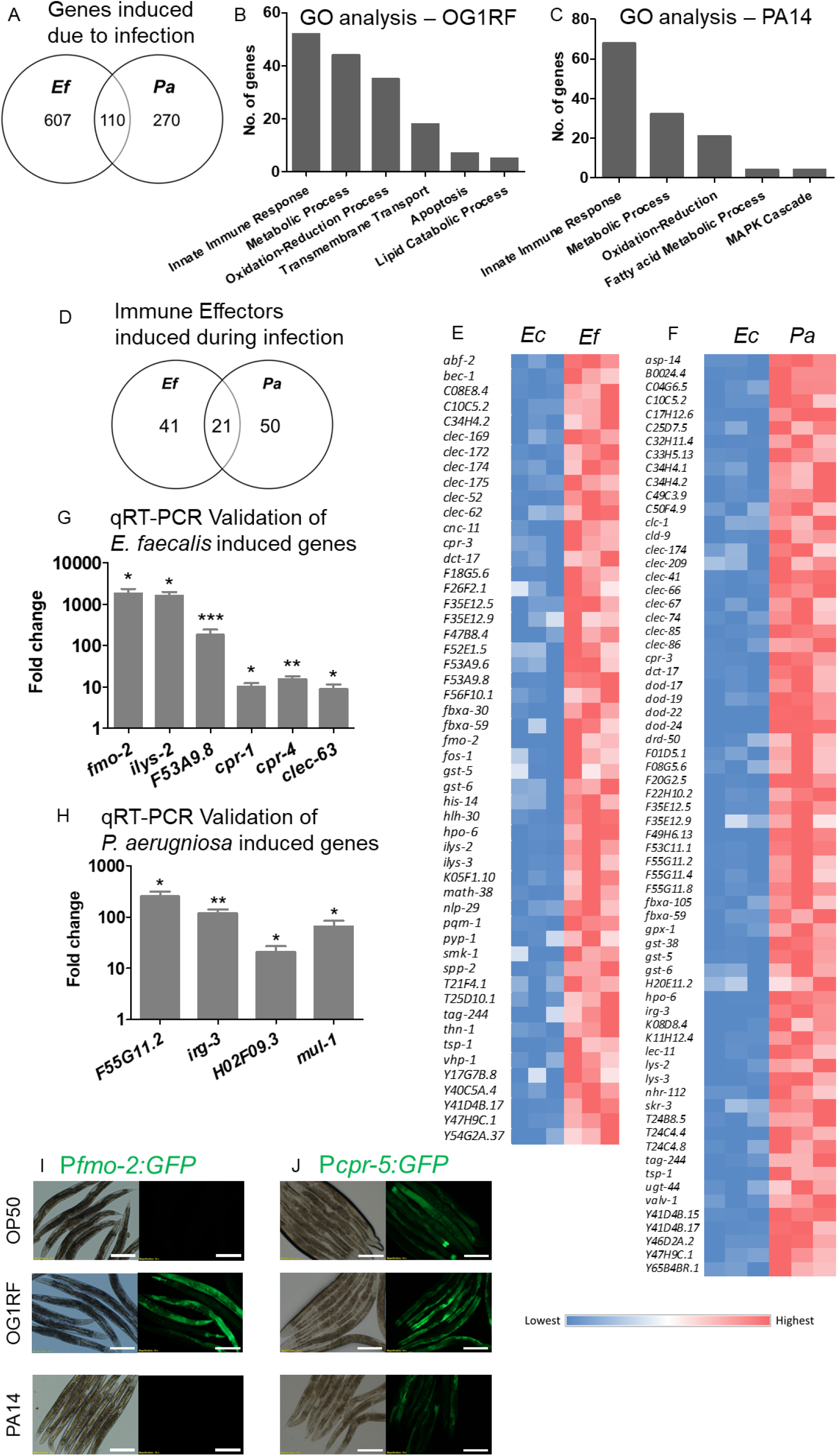
*C. elegans* early response to pathogens comprises of pathogen-specific and shared signatures. Venn diagram showing (A) genes induced 2-fold or higher (*p*≤0.05) upon exposure to *E. faecalis* (*Ef*) and *P. aeruginosa* (*Pa*), each compared to *E. coli* (*Ec*). (B) GO analysis of genes induced upon infection with *E. faecalis* in 8 hours. (C) GO analysis of genes induced upon infection with *P. aeruginosa* in 8 hours. (D) Innate immune response genes induced 2-fold or higher (*p*≤0.05) upon infection manifested in GO analysis. (E-F) Heat-map showing qualitative comparison in FPKM values for innate immune response genes in worms exposed to food (*E. coli*) and pathogen (E) *E. faecalis* and (F) *P. aeruginosa*. qPCR Validation of (G) *E. faecalis* induced immune effectors (H) *P. aeruginosa* induced immune effectors compared to food (*E. coli*). (I) P*fmo-2*:GFP transcriptional reporter upon exposure to *E. coli, E. faecalis* and *P. aeruginosa* (J) P*cpr-5*:GFP transcriptional reporter upon exposure to *E. coli, E. faecalis* and *P. aeruginosa*. Scale bar, 200 μm.

Gene ontology (GO) analysis of RNA seq data indicated that innate immune response, metabolic process, oxidation-reduction process, transmembrane transport, apoptosis, lipid catabolic process etc were enriched in the transcriptome of *C. elegans* exposed to *E. faecalis* and *P. aeruginosa* for 8 hours (Fig 1B, 1C). There were 52 and 68 innate immune response genes induced during *E. faecalis* exposure and *P. aeruginosa* exposure respectively (Fig 1E, 1G, see Table S1 and S2 for fold changes). Out of these, only 17 genes were shared between two exposures while the remaining were pathogen specific (Fig 1D). Signatures of immune response such as C-type lectins, aspartic proteases, cysteine proteases, glutathione transferases, and galectins were partly shared but most turned out to be pathogen-specific (Fig 1D; Supplementary Table S1 and S2) (Sun, Aballay and Singh, 2016). To further validate the specificity of this response, we utilized transcriptional reporters for *fmo-2*, encoding a inducible flavin mono oxygenase with detoxification function. On *E. coli, fmo-2* promoter had no basal activity, but it was induced strongly on *E. faecalis* in 8 hours while there was no appreciable induction on *P. aeruginosa* (Fig 1I). We also created a cysteine protease reporter strain, P*cpr-5*::GFP, and found that the reporter had a basal level expression on *E. coli* which was induced strongly on *E. faecalis* but not induced on *P. aeruginosa* (Fig 1J). We find that CPR-5 is expressed in *C. elegans* intestine which is the site of proliferation of *E. faecalis*. The pathogen-specific early response of *C. elegans* indicated that nematodes can differentiate between *P. aeruginosa* and *E. faecalis* and engage in a context specific response.

### Sensory input regulates pathogen-specific responses against *Enterococcus faecalis* and *P. aeruginosa*

The nervous system of *C. elegans* comprises over one thirds of the soma and enables the nematode to engage in robust chemotactic response to stimuli, learning and decision making. We hypothesized that ciliated neurons modulate pathogen specific immune effector signatures and survival on *E. faecalis* and *P. aeruginosa*. To address this, we utilized a mutation in *osm-6*, encoding a component of intraflagellar transport particle. The mutation affects sensory cilia of neurons due to and sensory perception (Bargmann, Hartwieg and Horvitz, 1993; Collet *et al*., 1998). We found that *osm-6(p811)* animals exhibited increased susceptibility to Gram positive bacterium *E. faecalis* compared to wild type N2 animals (Fig 2A). Wild type and *osm-6(p811)* animals had similar pumping rates on *E. faecalis* suggesting that their feeding was similar on pathogenic bacteria (Fig 2B). However, *E. faecalis* was better able to colonize the intestine of cilia mutant animals resulting in a higher CFU count per worm for *osm-6(p811)* animals than for WT animals (Fig 2C), suggesting deficient immune response in cilia mutants. Microscopic analysis of bacterial burden also showed higher *E. faecalis*::GFP bacteria in *osm-6(p811)* animals(Fig 2D,2E). We hypothesized that sensory perception of infection is required for optimal production of immune effector molecules. We looked at immune signatures composed of fifteen *E. faecalis* or *Staphylococcus aureus* infection induced genes utilizing our RNAseq analysis. We found that 14 out of 15 transcripts were upregulated in WT animals at 24 hours exposure to *E. faecalis* (Fig 2F–2M; S2A-S2G). Of these, four genes - *fmo-2*, aspartyl protease/*asp-10*, and *cpr-4 and cpr-5*, were not induced to wild type levels in *osm-6* cilia mutant animals (Fig 2F–2I). Transcripts for three lectin-like molecules, *clec-60, clec-62* and *clec-63*, were also not induced to wild type levels in *cilia* mutants during *E. faecalis* infection (Fig 2J–2L). F53A9.8 immune effector was upregulated >20 fold in WT animals by *E. faecalis* infection but induced <5 fold in *osm-6* animals. Six *E. faecalis* induced effector molecule (*cpr-1, cpr-8, clec-61, ech-9, ilys-2* and K05F1.10) appeared independent of OSM-6 sensory regulation (Fig S1A–S1G). In all, our analysis of 15 immune effectors indicated that a large fraction of *E. faecalis* induced immune effectors are regulated non-cell autonomously by ciliated neurons in *C. elegans*.

**Figure 2.**
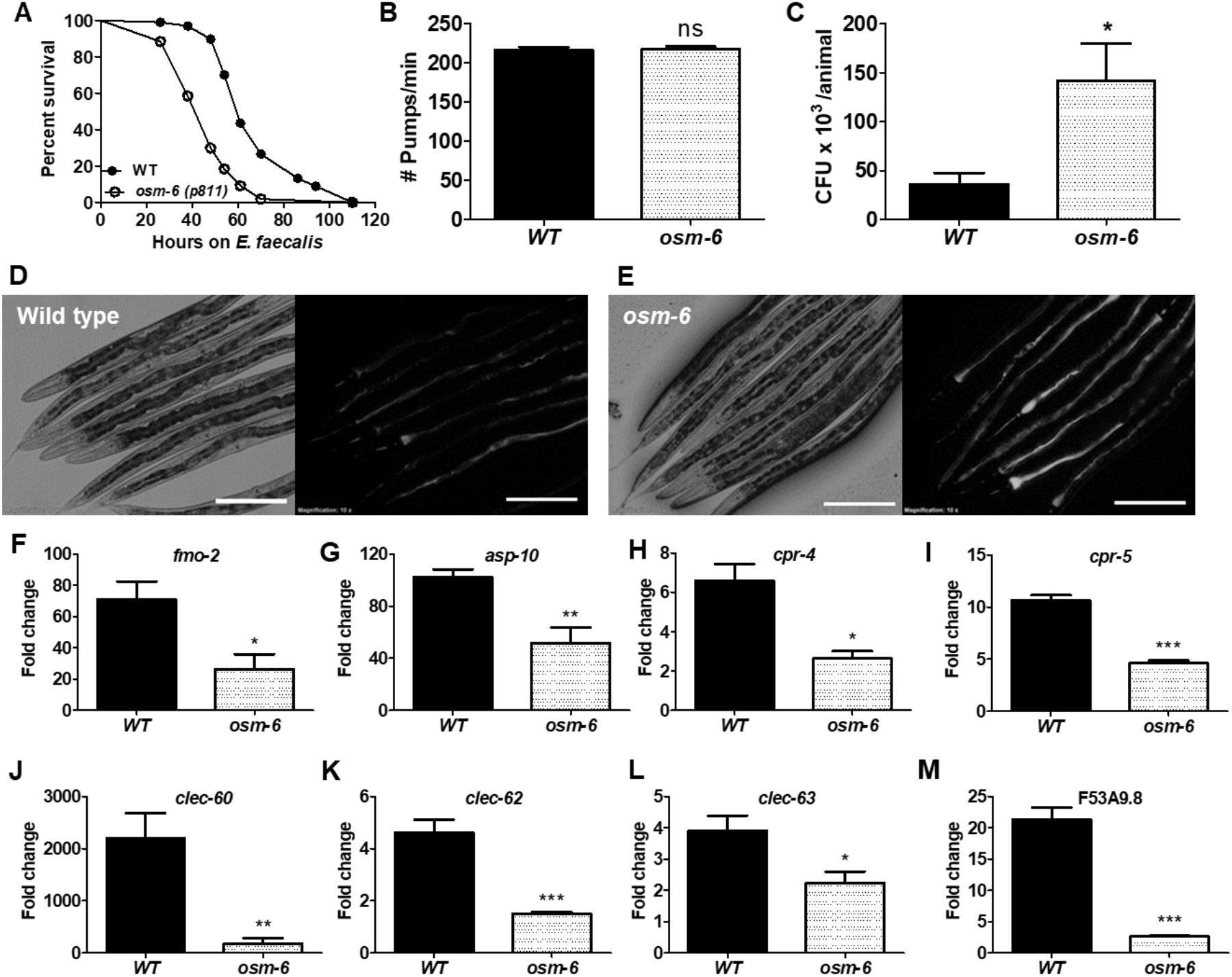
Sensory neurons regulate immunity to Gram positive bacteria *Enterococcus faecalis*. (A) Kaplan Maier survival curve for WT and *osm-6(p811)* animals exposed to *E. faecalis* (p<0.0001). (B) Pharyngeal pumping rates for WT and *osm-6(p811)* animals on *E. faecalis* at 4 hours of exposure. (C) Colony forming units recovered from WT and *osm-6(p811)* animals fed *E. faecalis*::GFP for 24 hours. Real time PCR analysis of immune effectors transcripts (F) *fmo-2*, Flavin monooxygenase, (G) *asp-10*, aspartic protease, (H) *cpr-4*, cysteine protease, (I) *cpr-5*, (J) *clec-60*, C type lectin, (K) *clec-62*, (L) *clec-63*, and (M) F53A9.8. Significance obtained by unpaired *t* test (ns > 0.05, * ≤0.05, **≤0.01, *** ≤0.001). Scale bar, 200 μm.

We found that cilia defective *osm-6(p811)* animals were highly susceptible to *P. aeruginosa* infection (Fig 3A) compared to WT animals. Consequently, we found that cilia mutants had higher bacterial burden in the intestine shown as CFU counts/animal (Fig 3C). To understand if ciliated neurons exert non cell autonomous control on immune effector expression in the intestine, the site of bacterial proliferation, we analysed 10 *P. aeruginosa* specific immune effectors for possible regulation by OSM-6. In the infection response gene family (Estes *et al*., 2010), *irg-1, irg-2* and *irg-3* induction was severely dampened in cilia mutants without any appreciable effect on *irg-7* induction (Fig 3D–3F, S2A) (Estes *et al*., 2010). Transcript levels of five other effectors-F55G11.2, *dod-24, mul-1*, F01D5.5 and C17H12.8-were also not induced to WT levels in *osm-6(p811)* animals (Fig 3G–3L) (Troemel *et al*., 2006; Powell, Kim and Ausubel, 2009). We found that a *P. aeruginosa* inducible lectin, *clec-67*, was also dependent on OSM-6 for induction. Thus, a large fraction (9 out of 10) of *P. aeruginosa* inducible effectors were modulated by OSM-6 dependent sensory perception in *C. elegans*. Taken together, analysis of *E. faecalis* and *P. aeruginosa* inducible effectors suggested that a large fraction of immune effectors repertoire is regulated non-cell autonomously by the nervous system of *C. elegans*. This also indicated that sensory perception is important for inducing immune response to pathogens.

**Figure 3.**
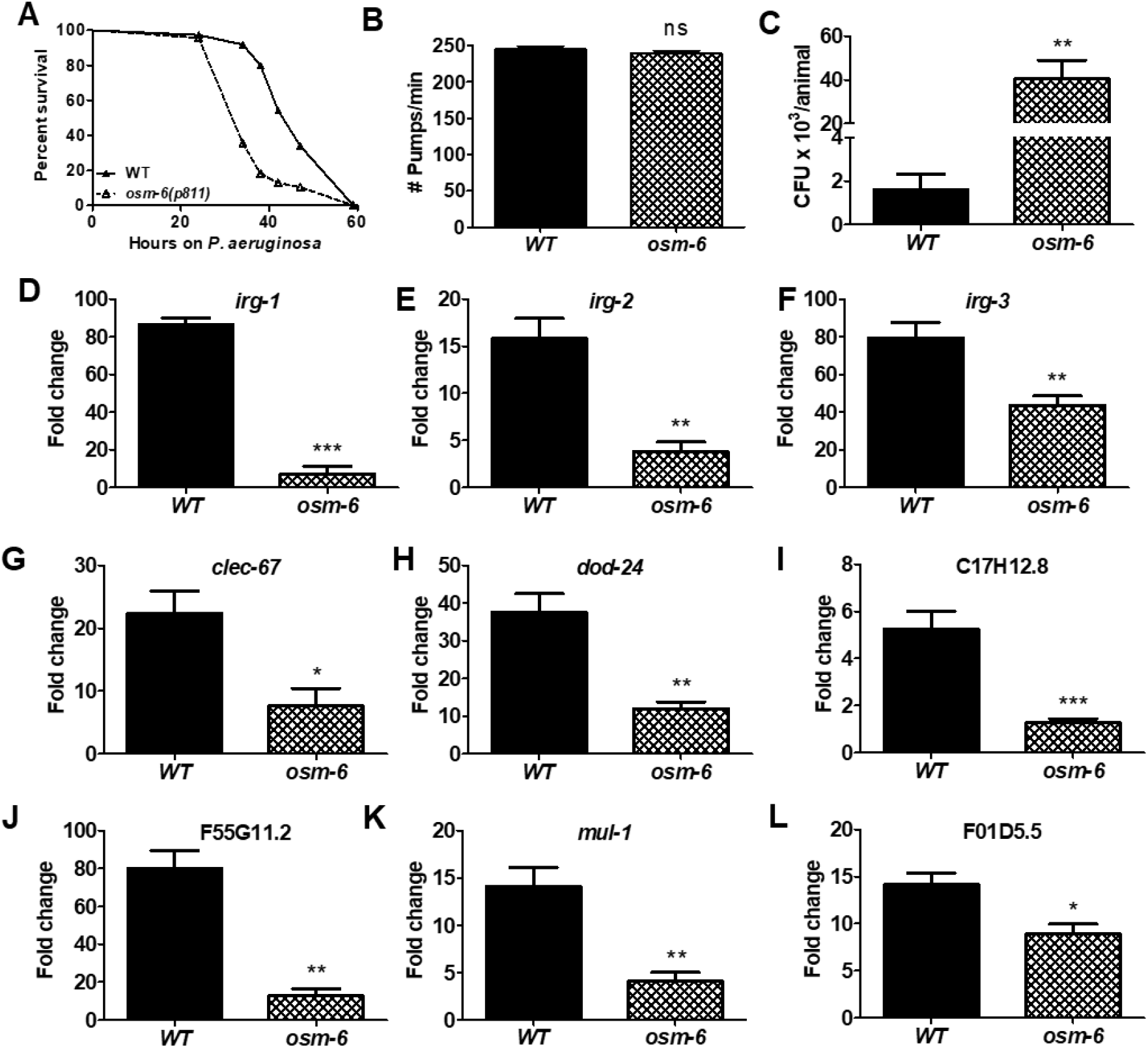
Sensory neurons regulate immunity to Gram negative bacterium *Pseudomonas aeruginosa*. (A) Kaplan Maier survival curve for WT and *osm-6(p811)* animals exposed to *P. aeruginosa* (p<0.0001). (B) Pharyngeal pumping rates for WT and *osm-6(p811)* animals on *P. aeruginosa* at 4 hours of exposure. (C) Colony forming units recovered from WT and *osm-6(p811)* animals fed *P. aeruginosa*::GFP for 24 hours. Real time PCR analysis of *P. aeruginosa* specific immune effectors transcripts (D) *irg-1*, infection response gene, (E) *irg-2*, (F) *irg-3*, (G) *clec-67*, C type lectin, (H) *dod-24*, (I) C17H12.8, (J) F55G11.2, (K) *mul-1* and (L) F01D5.5. Significance obtained by unpaired *t* test (ns > 0.05, * ≤0.05, **≤0.01, *** ≤0.001).

We were intrigued by the overwhelming reduction in both *E. faecalis* as well as *P. aeruginosa* inducible immune effector transcripts in cilia mutants. We questioned if the basal levels of these genes, on laboratory diet of *E. coli*, was controlled by the nervous system. When we compared the basal levels of 25 immune effector genes in *osm-6(p811)* and WT animals, we were surprised to see that while a number of them were expressed at the wild type levels, many immune effector transcripts were present at higher levels in *osm-6* animals than in the wild type (Fig S3A–S3B). This indicated that *osm-6(p811)* animals were not deficient in basal expression of immune effectors and suggested that neuronal sensing may be required for inducible expression of immune effectors.

### Neuroendocrine signalling regulates innate immune response of *C. elegans* to bacterial pathogens

*E. faecalis* and *P. aeruginosa* proliferate in the intestine of *C. elegans* (Tan, Mahajan-Miklos and Ausubel, 1999; Garsin *et al*., 2001). Consequently, majority of the immune effectors are expressed in the intestine (Estes *et al*., 2010; Irazoqui *et al*., 2010), the site of infection. Sensory perception of microbes by OSM-6 expressing neurons must be communicated to the intestine to modulate immune effector expression. Sensory neurons utilize neurotransmitters for synaptic transmission and neuropeptides for synaptic, extra synaptic, and medium to long range communications. Recent studies indicate a broad role for neuropeptide like proteins in control of physiological responses (Li, 2008; Nässel and Winther, 2010) as well as flight response to pathogenic bacteria (Cao *et al*., 2017; Lee and Mylonakis, 2017). Since intestine, the site of proliferation of *E. faecalis* or *P. aeruginosa*, is not innervated, neuropeptides present a possible mode of communication. We tested if disruption of neuropeptide processing due to mutation in proprotein convertase/EGL-3 could affect immune response to bacterial pathogens. We found that *egl-3(n150)* animals were susceptible to both *E. faecalis* and *P. aeruginosa* (data not shown. However, these animals had enhanced matricide linked to their egg laying defect. To correct for the *egl* phenotype, we treated *egl-3(n150)* animals with *cdc-25.1* RNAi to prevent germline proliferation and matricide and then analysed their susceptibility to pathogenic bacteria. We found that *egl-3(n150)*; *cdc-25.1* RNAi animals exhibited significant susceptibility towards *E. faecalis* but wild type susceptibility towards *P. aeruginosa* (Fig 4A, 4B) compared to WT; *cdc-25.1* RNAi on respective bacteria. The differential response neuropeptide processing deficient animals was really intriguing and prompted us to do further investigations. Pumping rates in both the WT; *cdc-25.1* RNAi and *egl-3(n150)*: *cdc-25.1* RNAi animals were similar, but colonization of the intestinal lumen was 8 times higher in *egl-3(n150)* animals than the wild type (Fig 4C–4E) indicating defective clearance of *E. faecalis* in these animals. Next, we checked immune response of *egl-3(n150)* animals to *E. faecalis*. We found that *egl-3(n150)* animals were defective in upregulating 3 proteases *−asp-10, cpr-5* and *cpr-8*. Other immune effectors that showed dysregulation in *osm-6(p811)* mutants did not show significant difference in *egl-3(n150)* mutants (Fig S4). In all, our data indicates that neuropeptide-based communication is necessary for modulating immune response to *E. faecalis*.

**Figure 4.**
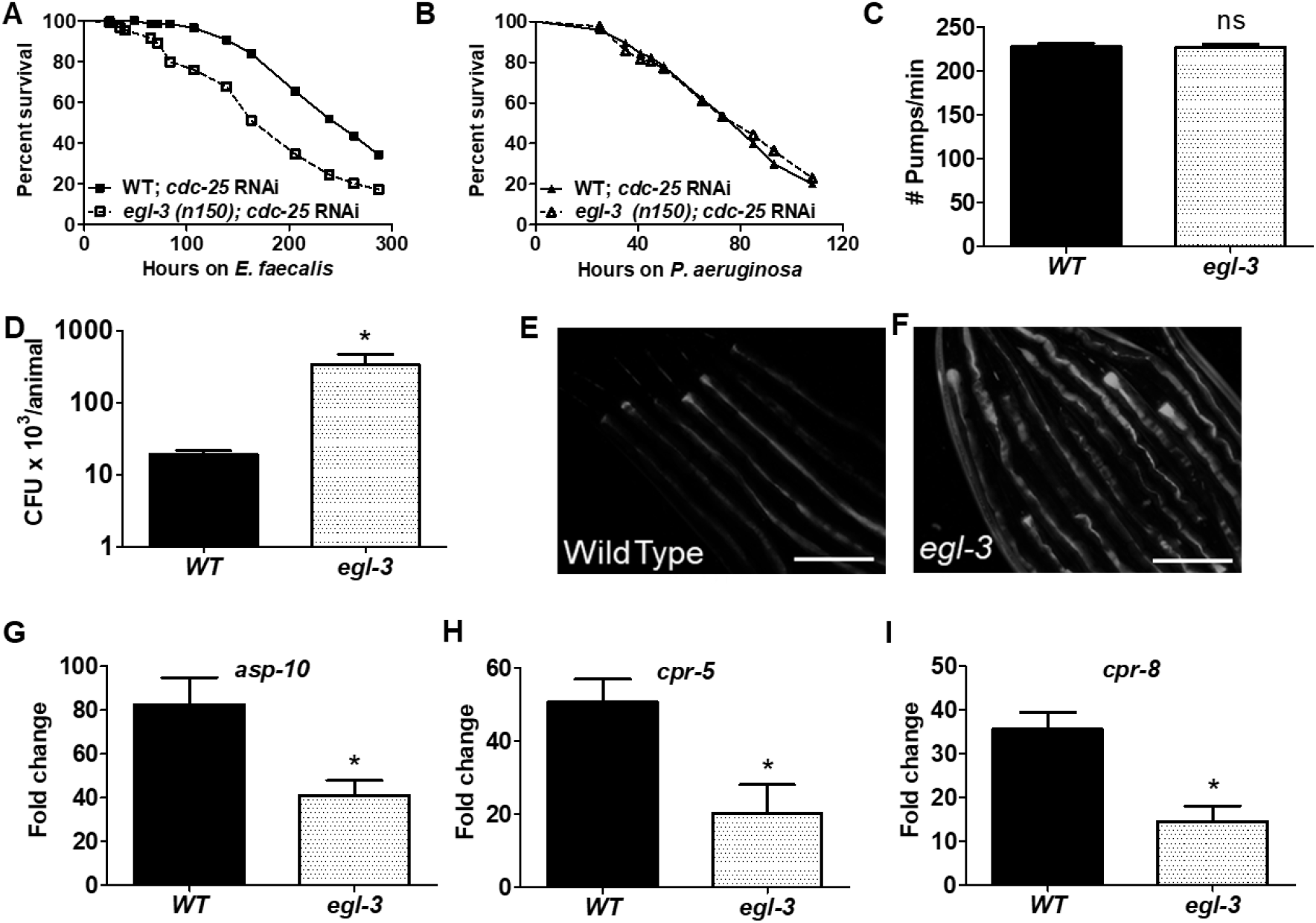
Neuropeptides differentially regulate survival and immune response to Gram negative and Gram positive bacteria. Kaplan Maier survival curve for WT and *egl-3(n150)* animals with *cdc-25.1* RNAi exposed to (A) *E. faecalis* (p<0.0001), and (B) *P. aeruginosa* (p, 0.62). (C) Pharyngeal pumping rates for WT; *cdc-25.1* and *egl-3(n150)*; *cdc-25.1 RNAi* animals on *P. aeruginosa* at 4 hours of exposure. (D) Colony forming units recovered from Wild-type and *egl-3(n150)*; *cdc-25.1 RNAi* animals fed *E. faecalis*::GFP for 24 hours. (E-F) Imaging of *E. faecalis*::GFP in (E) WT; *cdc-25.1* animals (F) *egl-3(n150)*; *cdc-25.1 RNAi* animals after 24 hours of exposure. Real time PCR analysis of *E. faecalis* specific immune effectors transcripts in WT; *cdc-25.1* and *egl-3(n150)*; *cdc-25.1 RNAi* animals, (G) *asp-10*, (H) *cpr-5*, and (I) *cpr-8*. Significance obtained by unpaired *t* test (ns > 0.05, * ≤0.05, **≤0.01, *** ≤0.001). Scale bar, 200 μm.

### OSM-6-FSHR-1 neuro-intestine axis regulates *C. elegans* resistance on *E. faecalis* as well as *P. aeruginosa*

To find mechanisms of immune effector regulation by ciliated neurons, we studied signalling pathways which act in *C. elegans* intestine. FSHR-1, an orthologue of human follicle stimulating hormone receptor, regulates immune homeostasis during infection as well as stress response in *C. elegans* intestine (Powell, Kim and Ausubel, 2009; Miller *et al*., 2015; Robinson and Powell, 2016; Yuen and Ausubel, 2018) independent of p38 MAP kinase or insulin signalling pathway. We asked if OSM-6 dependent sensory perception modulated FSHR-1 in the intestine upon exposure to pathogenic bacteria. First, we analysed the effect of *fshr-1* RNAi inhibition upon susceptibility in WT and in *osm-6(p811)* animals. As previously reported, RNAi inhibition of *fshr-1* in WT animals caused hypersusceptibility to *P. aeruginosa* (Fig 5A). However, RNAi inhibition of *fshr-1* in *osm-6(p811)* animals had significant but very small effect on their susceptibility compared to *osm-6(p811)*; vector control (Fig 5A). As shown in the mean survival (TD_50_) plot, *fshr-1* RNAi caused drastic (50%) decline in mean survival in WT animals but a more subtle (15%) decline in *osm-6* (*p811*) animals (Fig 5B) exposed to *P. aeruginosa*. We found that *fshr-1* RNAi in WT animals increased CFU load of *P. aeruginosa* (Fig 5C). In *osm-6(p811)* animals, CFU burden was higher in vector control and did not further increase due to *fshr-1* RNAi. This indicated that OSM-6-FSHR-1 axis regulates survival during *P. aeruginosa* infection. We reasoned that if OSM-6-FSHR-1 axis is important for resistance, these should regulate common set of immune effectors. As previously reported, FSHR-1 did not regulate immune response genes *−irg-1, irg-2, irg-3* and *irg-7* (Fig S5), but it regulated *clec-67*, F01D5.5, and F55G11.2-in WT animals exposed to *P. aeruginosa* (Fig 5D). However, *clec-67* and F01D5.5 levels were lower in *osm-6(p811)*; vector RNAi animals and did not reduce further due to *fshr-1* RNAi (Fig 5D–5E) indicating that OSM-6-FSHR-1 axis regulates expression of these *P. aeruginosa* specific immune effectors. Taken together, we were able to determine that OSM-6-FSHR-1 axis promoted *C. elegans* survival on *P. aeruginosa*, inhibited bacterial proliferation and led to upregulation of immune effectors during infection.

**Figure 5.**
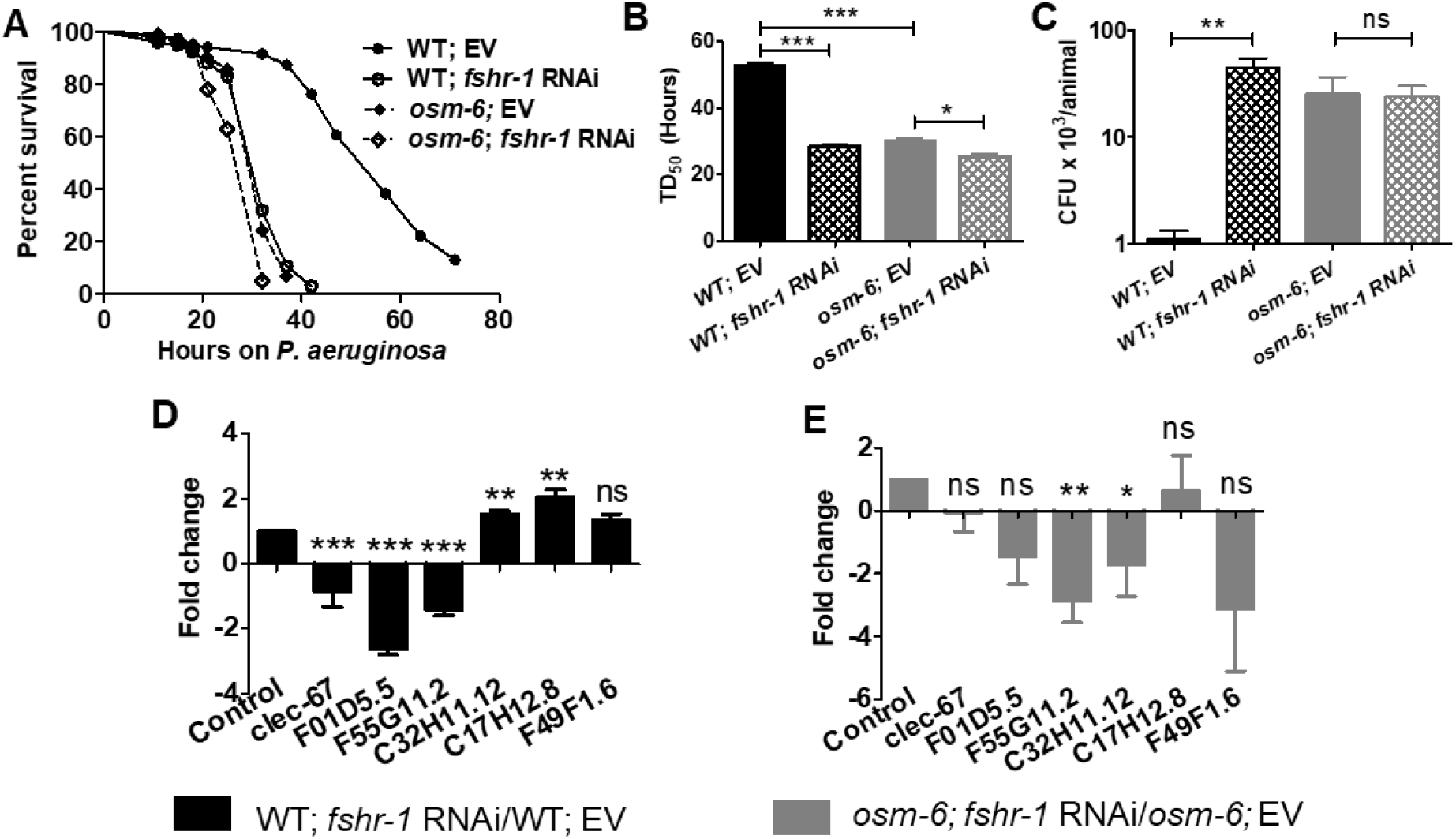
Sensory neurons regulate *fshr-1*-mediated immunity to Gram negative bacterium *P. aeruginosa*. (A) Kaplan Maier survival curve for WT and *osm-6(p811)* animals with vector and *fshr-1* RNAi on *P. aeruginosa*. (B) Mean survival (TD_50_) of WT and *osm-6(p811)* animals with vector and *fshr-1* RNAi. (C) Colony forming units of *P. aeruginosa*::GFP recovered from WT and *osm-6(p811)* animals with vector and *fshr-1* RNAi. Real time PCR analysis of *P. aeruginosa* specific immune effectors transcripts, (D) in WT animals grown on vector and *fshr-1* RNAi upon exposure to *P. aeruginosa* for 8 hours, (E) in *osm-6(p811)* animals grown on vector and *fshr-1* RNAi upon exposure to *P. aeruginosa* for 8 hours. Significance obtained by unpaired *t* test (ns > 0.05, * ≤0.05, **≤0.01, *** ≤0.001).

To test whether ciliated neurons regulate immunity via FSHR-1 signalling during *E. faecalis* infection as well, we performed epistasis analysis. We found that RNAi inhibition of *fshr-1* caused enhanced susceptibility to *E. faecalis* in WT animals, but less so in *osm-6(p811)* animals (Fig 6A–6B). RNAi inhibition of *fshr-1* also increased CFU load of *E. faecalis* in WT animals 10-fold compared to vector control, but not in *osm-6(p811)* animals (Fig 6C) suggesting that OSM-6 regulates immune resistance to *E. faecalis* infection via FSHR-1 pathway. We checked the *E. faecalis* mediated inducibility of OSM-6 dependent immune effectors in WT animals with *fshr-1* RNAi over vector RNAi and found that *fmo-2, clec-62, clec-63* and F53A9.8 were not regulated by FSHR-1 (Fig 6D–6E, S6A-B). However, we found that *fshr-1* RNAi suppressed pathogen induced upregulation of *cpr-4, cpr-5 and asp-10* in WT animals (Fig 6D) showing for the first time that specific *E. faecalis* effectors are regulated by FSHR-1 pathway. Interestingly, *fshr-1* RNAi did not suppress *cpr-4, cpr-5 and asp-10* in *osm-6(p811)* animals (Fig 6E) indicating that *cpr-4, cpr-5* and *asp-10* are regulated by OSM-6-FSHR-1 axis. In all, our data provides evidence for non-cell autonomous regulation of FSHR-1 by the nervous system, in regulating differential immune response to *E. faecalis* and *P. aeruginosa*.

**Figure 6.**
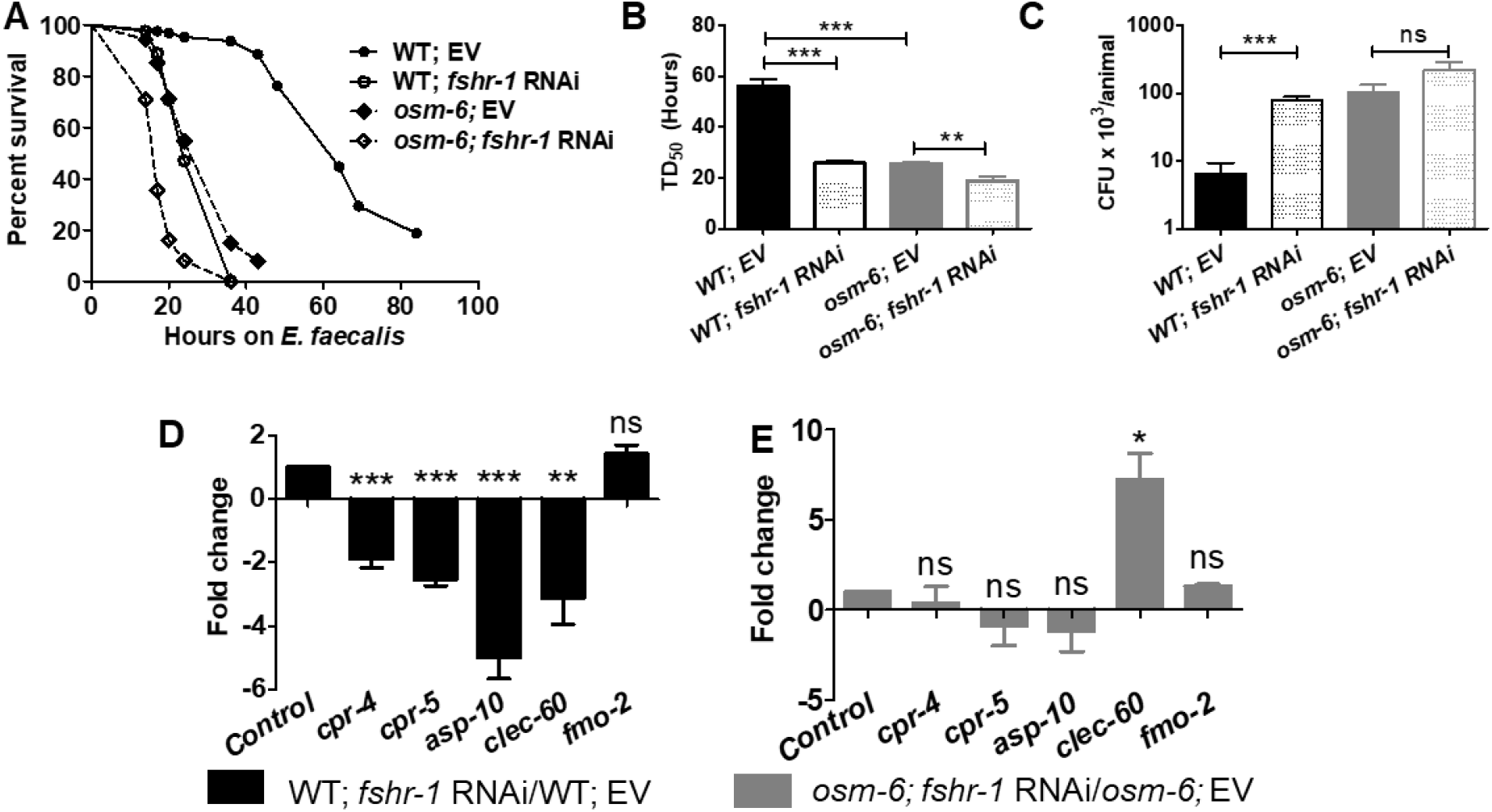
Sensory neurons regulate *fshr-1*-mediated immunity to Gram positive bacterium *E. faecalis*. A) Kaplan Maier survival curve for WT and *osm-6(p811)* animals with vector and *fshr-1* RNAi on *E. faecalis*. (B) Mean survival (TD_50_) of WT and *osm-6(p811)* animals grown with Vector and *fshr-1* RNAi. (C) Colony forming units of *E. faecalis*::GFP recovered from WT and *osm-6(p811)* animals with Vector and *fshr-1* RNAi. Real time PCR analysis of *E. faecalis* specific immune effectors transcripts, (D) in WT animals grown on vector and *fshr-1* RNAi upon exposure to *E. faecalis* for 8 hours, (E) in *osm-6(p811)* animals grown on vector and *fshr-1* RNAi upon exposure to *E. faecalis* for 8 hours. Significance obtained by unpaired *t* test (ns > 0.05, * ≤0.05, **≤0.01, *** ≤0.001).

### OSM-6-HLH-30-FMO-2 neuro-intestine axis controls *C. elegans* immune tolerance to *E. faecalis*

Our analysis of OSM-6-FSHR-1 axis indicated that although this axis regulated survival during infection, it did not control all OSM-6 regulated immune effectors pointing to existence of additional brain-gut axes of immunity. We found that *fmo-2*, induced >1000-fold during *E. faecalis* exposure (Fig 1G, Fig1I), was not dependent on FSHR-1. However, RNAi knockdown of *fmo-2* in WT animals caused enhanced susceptibility to *E. faecalis* infection (Fig 7A) suggesting that FMO-2 activity promotes survival on *E. faecalis*. On the other hand, RNAi inhibition of *fmo-2* in *osm-6(p811)* animals did not cause enhanced susceptibility (Fig 7A) as expected since *E. faecalis* induced *fmo-2* transcripts were lower in *osm-6* cilia mutant (Fig 2F). FMO-2 function in the intestine improves health span, induces proteostasis, and it is dependent on helix-loop-helix transcription factor, HLH-30 (Rourke and Ruvkun, 2014; Leiser *et al*., 2015). Our RNA-seq data showed that *hlh-30* itself was induced 2-fold upon *E. Faecalis* infection (Fig 1E, Table S1) making it one of a few infection inducible transcription factors (Table S1). By qRT-PCR analysis of *E. faecalis* exposed animals, we found that *hlh-30* transcripts were indeed upregulated in WT animals but not in *osm-6(p811)* animals (Fig 7B). We found that *hlh-30* RNAi in WT animals caused enhanced susceptibility to *E. faecalis* (Fig 7C-D) however, *hlh-30* RNAi had no effect on the survival of *osm-6 (p811)* animals (Fig 7C-D) suggesting that an OSM-6-HLH-30 axis of immune response is active during *E. faecalis* infection of *C. elegans*.

**Figure 7.**
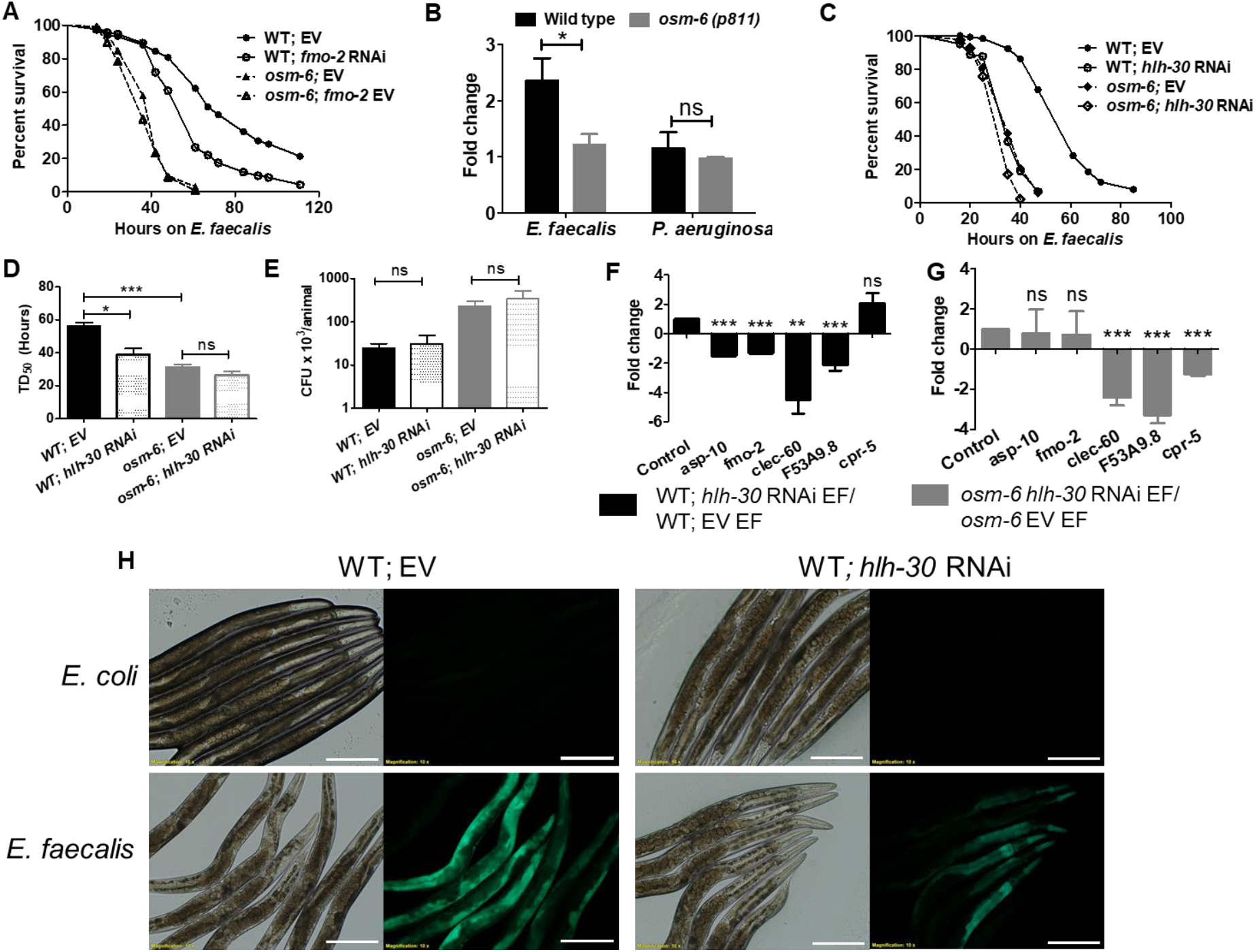
Sensory neurons regulate HLH-30 mediated immunity to Gram positive bacterium *E. faecalis*. (A) Kaplan Maier survival curve for WT and *osm-6(p811)* animals with vector and *fmo-2* RNAi on *E. faecalis*. (B) Real Time PCR analysis of *hlh-30* transcript levels in WT and *osm-6(p811)* animals upon exposure to *E. faecalis* and *P. aeruginosa*. (C) Kaplan Maier survival curve for WT and *osm-6(p811)* animals with vector and *hlh-30* RNAi on *E. faecalis*. (D) Mean survival (TD_50_) of WT and *osm-6(p811)* animals grown on vector and *hlh-30* RNAi followed by exposure to *E. faecalis*. (E) *E. faecalis*::GFP colony forming units recovered from WT and *osm-6(p811)* animals grown on vector and *hlh-30* RNAi upon 18 hours exposure. (F-G) Real time PCR analysis of *E. faecalis* specific immune effectors transcripts in (F) WT animals grown on vector and *hlh-30* RNAi and exposed to *E. faecalis* for 8 hours, (G) *osm-6(p811)* animals grown on vector and *hlh-30* RNAi and exposed to *E. faecalis* for 8 hours. (H) Effect of *hlh-30* RNAi on P*fmo-2*::GFP expression upon exposure to *E. coli* and *E. faecalis*. Significance obtained by unpaired *t* test (ns > 0.05, * ≤0.05, **≤0.01, *** ≤0.001). Scale bar, 200 μm.

To understand how OSM-6-HLH-30 axis might afford protection to *C. elegans* during *E. faecalis* infection, we analysed expression of pathogen specific immune effectors. We found that *E. faecalis* mediated induction of *fmo-2* transcript was lower in WT; *hlh-30* RNAi animals compared to WT; vector animals (Fig 7F). *E. faecalis* induced fmo-2 transcription was not altered in *osm-6(p811)* animals by *hlh-30* RNAi (Fig 7G). compared to vector control animals also exposed to *E. faecalis*. Using the P*fmo-2*::GFP reporter, we found that *E. faecalis* induced *fmo-2* was dependent partly on HLH-30 (Fig 7H). Additional *E. faecalis* induced immune effectors-*asp-10, clec-60* and F53A9.8-were also dependent on HLH-30 (7F-G). Of these, *asp-10* was regulated by OSM-6-HLH-30 axis (Fig 7F-G, S7C-D). Despite HLH-30 requirement of survival and immune effector expression during *E. faecalis* infection, it did not regulate *E. faecalis* proliferation in *C. elegans. hlh-30* RNAi caused no increase in CFU burden in *C. elegans* intestine, compared to respective vector RNAi controls (Fig 7E). This suggested that HLH-30 imparts tolerance to *E. faecalis* infection. In all, our results show that OSM-6-HLH-30 neuro-intestine axis regulates immune tolerance to *E. faecalis* via its effect on *fmo-2* and *asp-10* transcripts.

RNAseq and reporter expression data showed that *fmo-2* is not induced upon *P. aeruginosa* exposure (Table S2, Fig 1I). Even, *hlh-30* was not induced upon exposure to *P. aeruginosa* (Fig 1E, 7B) and *hlh-30* RNAi in WT animals caused slight albeit significant increase in susceptibility to *P. aeruginosa* (Fig S7A, S7B) suggesting that this transcription factor has minor role in response to Gram negative bacterium.

We describe a model of immune regulation by *C. elgans* ciliated neurons (Fig 8) wherein OSM-6 expressing ciliated, sensory neurons regulate FSHR-1/GPCR and HLH-30/TFEB transcription factor during exposure to pathogenic bacteria. OSM-6-FSHR-1 axis regulates nematode’s survival on *P. aeruginosa* and synthesis of at least two specific immune effectors-*clec-67* and F01D5.5. OSM-6-FSHR-1 axis regulates resistance to *E. faecalis* infection as well, via its effect on immune effectors-cpr-4, *cpr-5*, and *asp-10*. An additional sensory neuron-intestine axis, OSM-6/HLH-30 axis-regulates tolerance to *E. faecalis* infection via its effect on a subset of *E. faecalis* induced immune effectors-*asp-10* and *fmo-2*. Altogether, we show that sensory neurons modulate at least two different immunity axes in *C. elegans* intestine to execute pathogen tailored immune response to microbes.

**Figure 8.**
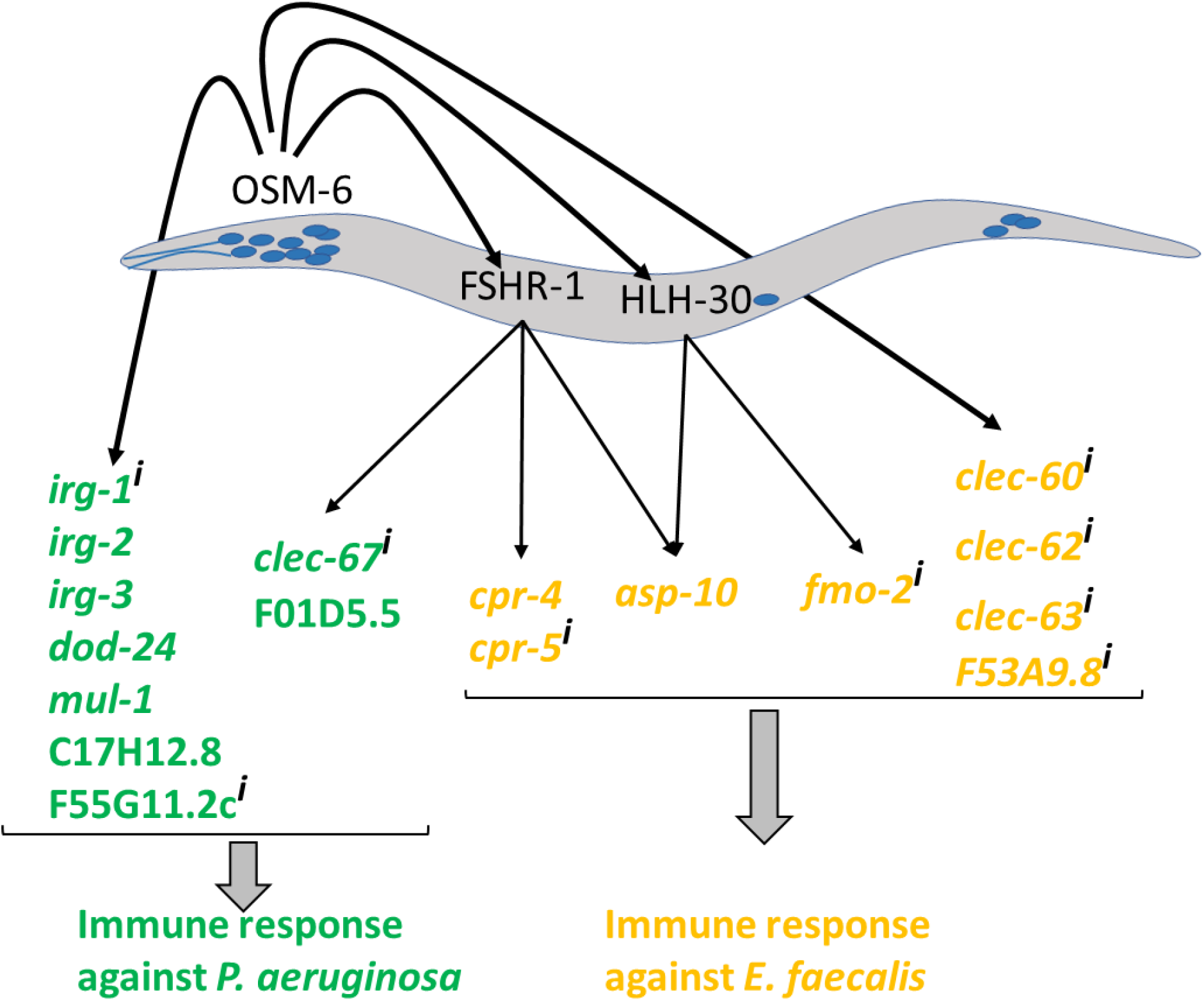
Sensory neurons regulate immune response via FSHR-1/GPCR and HLH-30/TFEB in the intestine. OSM-6 dependent ciliated neurons of *C. elegans* enable the nematode to elicit immune response in non-cell autonomous manner via FSHR-1/GPCR and HLH-30/TFEB in the intestine. FSHR-1 regulates pathogen specific immune effectors to impart resistance to *P. aeruginosa* and *E. faecalis*. HLH-30 provides tolerance to *E. faecalis* infection. *P. aeruginosa* induced effectors are marked in green, and *E. faecalis* induced effectors in yellow. Immune effectors with expression in the intestine are denoted with ^i^.

## DISCUSSION

In this study, we asked whether sensory neurons of *C. elegans* allow it to differentiate between Gram positive and Gram negative bacteria in order to mount tailored immune responses. We show that transcription response of *C. elegans* to *E. faecalis* and *P. aeruginosa* involves ~75% of non-overlapping immune signatures (Fig 1). Sensory perception by ciliated neurons regulates optimal production of 8/15 of *E. faecalis* specific immune effectors tested and 9/10 of *P. aeruginosa* specific immune effectors tested. Ciliated neurons utilize two different signalling mechanism in the intestine to modulate immune effector expression and survival on *E. faecalis*. One of these neuro-intestine axes also mediates survival on *P. aeruginosa*. Thus, we provide evidence for sensory control of differential immune response to Gram negative and Gram positive bacteria.

*C. elegans* nervous system is composed of 302 neurons of which 60 are ciliated sensory neurons. These include amphid sensory neurons in the head, labial neurons, mechanosensory neurons and O_2_/CO_2_ sensing neurons in the pseudo coelomic cavity (Scholey, 2007). Although, *osm-6* mutant animals were shown to have enhanced lifespan on *E. coli* (Apfeld and Kenyon, 1999), their enhanced susceptibility to pathogenic bacteria is very interesting. This enhanced susceptibly is due to the inability of these mutants to activate two pathways involved in immune homeostasis. Ciliated neurons are required for optimal induction of at least 8 *E. faecalis* specific effectors and at least 9 *P. aeruginosa* specific immune effectors. 10 of these immune effectors have expression in the intestine underscoring the importance of non-cell autonomous control. We also observed a high basal level of immune effectors in *osm-6 (p811)* animals. This indicates that sensory neurons might participate in a pathogen surveillance program and inhibit basal expression of immune effectors when they are not needed.

We were surprised by presence of HLH-30 dependent survival mechanism in *E. faecalis* exposed animals, but not in *P. aeruginosa* infected animals. We found that HLH-30 was transcriptionally upregulated upon *E. faecalis* exposure in OSM-6 dependent manner, but not regulated by *P. aeruginosa* infection. HLH-30 also mediates *C. elegans* survival on *Staphylococcus aureus* infection by regulating a battery of immune effectors and cytoprotective genes, in addition to components of autophagy pathway (Visvikis *et al*., 2014). Although there was no overlap between HLH-30 controlled genes in *E. faecalis* infection in this study (Figure 7) and in *S. aureus* infection (3), both bacteria are Gram positive cocci with similar features raising an interesting possibility that ciliated neurons might also regulate HLH-30 activity during *S. aureus* infection.

*C. elegans* response to *E. faecalis* has been shown to involve production of reactive oxygen species by the host for elimination of pathogenic cocci (Garsin *et al*., 2001; Chávez *et al*., 2007; Mohri-Shiomi and Garsin, 2008; Chávez, Mohri-Shiomi and Garsin, 2009; van der Hoeven *et al*., 2011; Tiller and Garsin, 2014; McCallum and Garsin, 2016; Swoboda *et al*., 2016; Liu *et al*., 2019). *C. elegans* also upregulates a number of detoxification enzymes involved in oxidative stress response (van der Hoeven *et al*., 2011; McCallum and Garsin, 2016) suggesting that this is likely a tolerance mechanism of survival. ROS has not been looked at in *P. aeruginosa* infection of nematodes. In the scenario where ROS production by *C. elegans* is not an active mechanism of antibacterial response during *P. aeruginosa* infection, it is conceivable that a certain arm of tolerance mechanism may not be necessary. This might explain very mild susceptibility phenotype of *hlh-30* RNAi animals during *P. aeruginosa* infection.

We have earlier shown that non cell autonomous control of immunity by the nervous system, during *P. aeruginosa* infection, impinges on p38 MAPK pathway and non-canonical UPR pathways (Sun *et al*., 2011; Cao *et al*., 2017). C. elegans has approximately 120 cholinergic neurons and these can activate *wnt* signalling in the intestine in response to infection by *S. aureus* (Irazoqui *et al*., 2008; Labed *et al*., 2018). Our study showcases the involvement of two ciliated neuron-intestine axes in this nematode to facilitate differential response to pathogenic microbes. OSM-6-FSHR-1 axis facilitates tailored response to *P. aeruginosa* and *E. faecalis* while OSM-6-HLH-30 axis appears essential for nematode’s response to *E. faecalis*. The latter is an example of a pathogen specific neuro intestine axis of immune tolerance. FSHR-1 and HLH-30 do not control all expression of some OSM-6 dependent immune effectors such as *clec-62, clec-63* and F53A9.8 suggesting that other signalling mechanisms in the intestine must regulate them. OSM-6 regulated immune effectors CLEC-60 is a C type lectin. It is under the control of wnt pathway in *S. aureus* infection (Labed *et al*., 2018). Wnt pathway function in *E. faecalis* infection and its control by ciliated neurons would be an interesting question to be pursued in future. We show that ciliated neurons are required for upregulation of *P. aeruginosa* specific effectors such as infection regulated gene or *irg* family, earlier reported to be regulated by bZIP transcription factor ZIP-2 (Estes *et al*., 2010). However, RNAi inhibition of *zip-2* did not cause susceptibility to *P. aeruginosa* infection in WT or cilia mutants (data not shown) suggesting that this axis does not regulate survival. Additional *P. aeruginosa* responsive immune effectors-C17H12.8, F55G11.2, *mul-1* are under the control of p38 MAPK in *P. aeruginosa* infection (Troemel *et al*., 2006). We have earlier shown that NPR-1 expressing neurons, also ciliated, positively regulate p38 MAPK activity (Styer *et al*., 2008). This suggests that ciliated neurons likely activate a number of signalling mechanisms during infection in *C. elegans* in a context (pathogen) specific manner.

An interesting area of research, leading from our study, would be identification of individual or subsets of ciliated neurons that regulate survival on Gram positive and Gram negative bacteria. Our previous work with GPCR adaptor protein Arr-1 in the nervous system indicated that while ciliated neurons – AFD, ADF, ASH/ASI, and AQR/PQR/URX-regulated survival on *P. aeruginosa* and *Salmonella enterica* (Singh and Aballay, 2012), these neurons did not have an impact upon *C. elegans* survival on *E. faecalis*. Analysis of control of *P. aeruginosa* and *E. faecalis* inducible early response immune signatures by individual neurons will help in deciphering pathogen specific response of nematodes. Systematic ablation of ciliated neurons would be instrumental in understanding of how sensory perception modulates resistance and tolerance to infection.

Conventionally, professional immune cells harbouring membrane localised or cytosolic pattern recognition receptors are believed to recognise microbe-associated molecular patterns to control innate immune response. Discovery of G protein coupled receptors on mouse nociceptive neurons for sensing *S. aureus* formylated peptides or MAMP has changed this notion (Chiu *et al*., 2013; Pavlov and Tracey, 2017) and has paved the way for pharmacological management for pain independent of involvement of neutrophils and cytokines (Adams *et al*., 2017). Our study provides a parallel for immune activation in *C. elegans* and suggests broader implications for study of FSHR and TFEB pathway in inflammation and immunity.

## METHODS

### Bacterial Strains

Bacterial strains *Escherichia coli* OP50, *Enterococcus faecalis* OG1RF, and *Pseudomonas aeruginosa* PA14 were used (Brenner, 1974; Singh and Aballay, 2006).

### *C. elegans* strains

*C. elegans* Bristol N2 (wild type); *osm-6* (p811) *egl-3* (n150) and seaEx13 [*fmo*-2p::GFP + unc-119(+)] were used.

### Generation of cpr-5p::GFP transgenic strain

A 644-bp genomic fragment containing *cpr-5* promoter was amplified using the following primers – 5’ ttggcatgcctttcttatcattgatttgaag3’ and 3’ttgggatcctatgagagaagtgtctgcg5’ and cloned into the SphI and BAMHI sites of pPD95_77 vector creating a transcriptional fusion. Wild type animals were microinjected with 5ng/μl of construct to generate VSL1903, agEx(*cpr*-5p::GFP + *unc*-122p::dsRed). The plasmid was maintained as extrachromosomal array.

### Survival Assay

*C. elegans* strains were maintained at 20°C on NGM agar plates seeded with *E. coli* OP50. *E. faecalis* cultures were grown in Brain-Heart Infusion (BHI) broth at 37°C for 5 hours and bacterial lawns were prepared by spreading 50 μl of culture on BHI Agar plates. *P. aeruginosa* cultures were grown in Luria-Bertani at 37°C overnight and bacterial lawns were prepared by spreading 50 μl of culture on modified NGM (Slow-killing) plates. Plates were incubated at 37°C before seeding then with young adult worms grown at 20°C. Assays were performed at 25°C and animals were scored for death and transferred every day to fresh plates. Animals were considered dead if they failed to respond to touch. All survival assays were done at least 3 times with more than 80 worms in each replicate.

### RNA interference

*C. elegans* were allowed to lay eggs and develop on *E. Coli* HT115 bacteria expressing double stranded RNA against target genes. Young adults grown on RNAi bacteria were used to from egg to young adult stage.

### Quantification of bacterial load

Young adult worms were allowed to feed on PA14::GFP or OG1RF::GFP for 24 hours at 25°C. Three to four replicates of 10 animals each were washed three times with M9 buffer, then crushed to release bacterial cells. Serial dilutions were plated to count CFU/worm using the following formula-

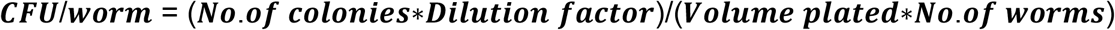

### RNA Isolation and qPCR

Young adult worms were exposed to bacteria were harvested in Qiazol reagent. RNA was isolated using Qiagen Universal RNA isolation kit. Genomic DNA was removed using Thermo Fisher DNAse I. cDNA was prepared using BIORAD iScript cDNA Synthesis Kit. qPCR was conducted using BIORAD iTAQ Universal SYBR Green Supermix on an Applied Biosystems Quant Studio 3 machine. C_⊤_ values obtained were used to calculate fold change using Livak method (Schmittgen and Livak, 2008). All target genes were normalized to act-1/Actin. Primer sequences are available upon request.

### Fluorescence Microscopy

Adult *fmo-2*P::GFP and adult *cpr-5*P::GFP animals were allowed to feed on OP50, OG1RF and PA14 for above mentioned time. 10-12 worms were mounted on fresh agarose-pad slides and imaged using fluorescence microscope Olympus IX81.

### RNA-sequencing and Data Analysis

RNA was isolated from L4 worms, fed *E. coli* OP50, *P. aeruginosa* PA14 or *E. faecalis* OG1RF for 8 hours, using Qiagen Universal RNA isolation kit (three biological replicates). cDNA library was prepared using NEBNext Ultra II Directional RNA Library Prep Kit from Illumina. The samples were then subjected to 50-base pair single-end sequencing resulting in 20 million reads using Illumina sequencer at Genotypic Pvt Ltd. The raw data was analysed using a pipeline described recently (Pertea *et al*., 2016). HISAT was used to map the RNA-seq reads onto the latest *C. elegans* reference genome (WS266) with output in the form of Sequence Alignment Map (SAM). HISAT was able to map ~95%of the genome. SAM format was converted to its binary form BAM using SAMTOOLS. BAM files were used as input for assembling the transcripts using STRINGTIE. Stringtie was used to assemble and quantify the levels of expressed genes to produce Fragments Per Kilobase of exon per Million fragments mapped (FPKM) values. The assembly was merged into a singular Gene Transfer Format (GTF) to facilitate comparison with the reference annotation in the same format. Fold change in expression for was calculated using the following formula-

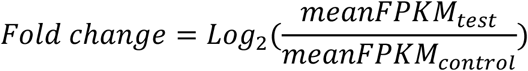

*E. faecalis* or *P. aeruginosa* upregulated genes were shortlisted with a cut off of 2-fold or above, and P-values less than 0.05. Raw RNAseq data has been deposited in the NCBI BioProject database (accession number PRJNA521879, and PRJNA524750).

### Gene Ontology Analysis

Genes with a cut off of 2-fold or above, and P-values less than 0.05 were considered for Gene Ontology enrichment analysis using the DAVID tool. Unique wormbase IDs were used as input data.

### Statistical Analyses

Survival plots were plotted as Kaplan Meier curves using the GraphPAD PRIZM and Logrank test was used to compare survival curves. Survival curves were considered different if p-value < 0.05. Unpaired t-test was used to analyse CFU load, Pharyngeal pumping and qRT-PCR results. P-values ≤0.05 was considered significant. Star annotations - ns > 0.05, * ≤0.05, **≤0.01, *** ≤0.001.

## ACKNOLEDGMENTS

We thank Prof Manuel Espinosa, Spanish National Research Council for *Enterococcus faecalis* strains. Some *C. elegans* strains were provided the CGC which is funded by the NIH Office of Infrastructure Programs (P40 OD01440). We thank Sandhya Visveswaraiah and Sambuddho Mukherjee for critical reading of the manuscript. We thank Sanchit Garg and Divakar Badal for providing technical support towards RNA-seq data analysis. This work was supported by the Wellcome Trust/DBT India Alliance Intermediate Fellowship (Grant no. IA/I/13/1/500919) awarded to Varsha Singh.

## AUTHOR CONTRIBUTIONS

AG, MMV and VS conceptualized the project. AG performed the experiments. VS, AG and MMV analysed the data and interpreted the results. VS, AG and MMV wrote the manuscript.

---

**Supplementary Figure S1.**
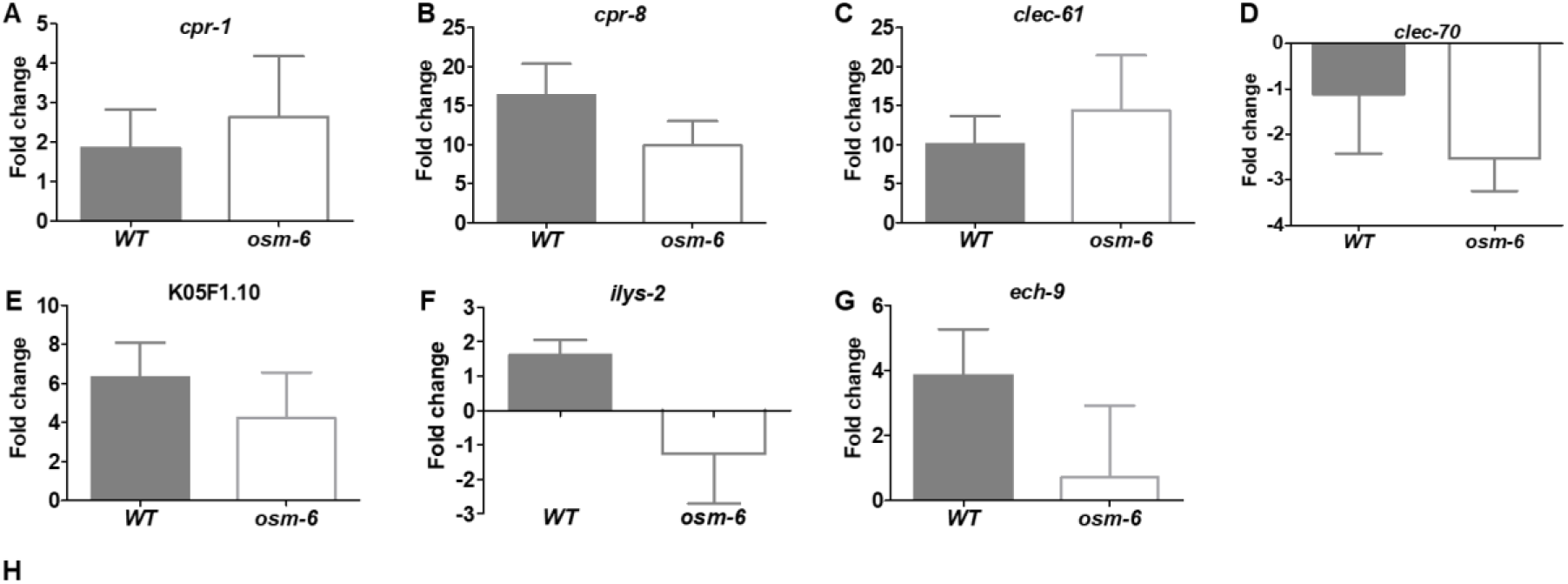
Real time PCR analysis of immune effectors in WT and *osm-6(p811)* animals exposed to *E. faecalis*. (A) *cpr-1*, cysteine protease (B) *cpr-8*, cysteine protease, (C) *clec-61*, C-Type Lectin, (D) *clec-70*, C-Type Lectin, (E) K05F1.10, hypothetical gene, (F) *ilys-2*, invertebrate lysozyme, and (G) *ech-9*, Enoyl-CoA Hydratase. Significance obtained by unpaired *t* test (ns > 0.05, * ≤0.05, **≤0.01, *** ≤0.001).

**Supplementary Figure S2.**
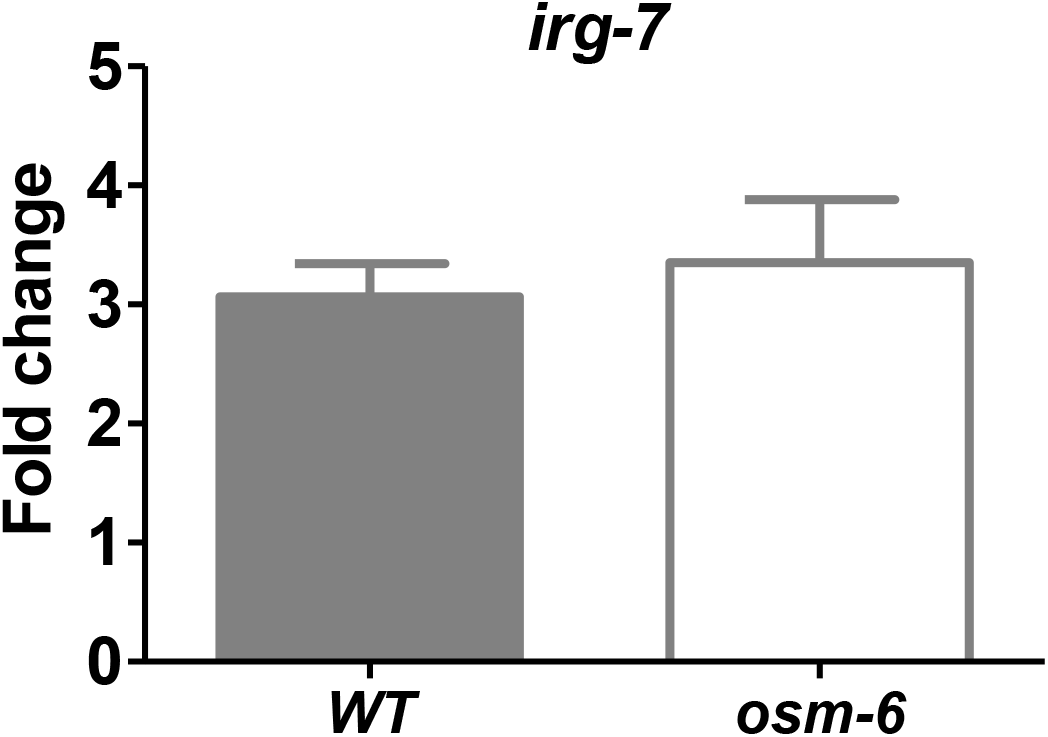
Real time PCR analysis of immune effectors transcript *irg-7*, in WT and *osm-6(p811)* animals.

**Supplementary Figure S3.**
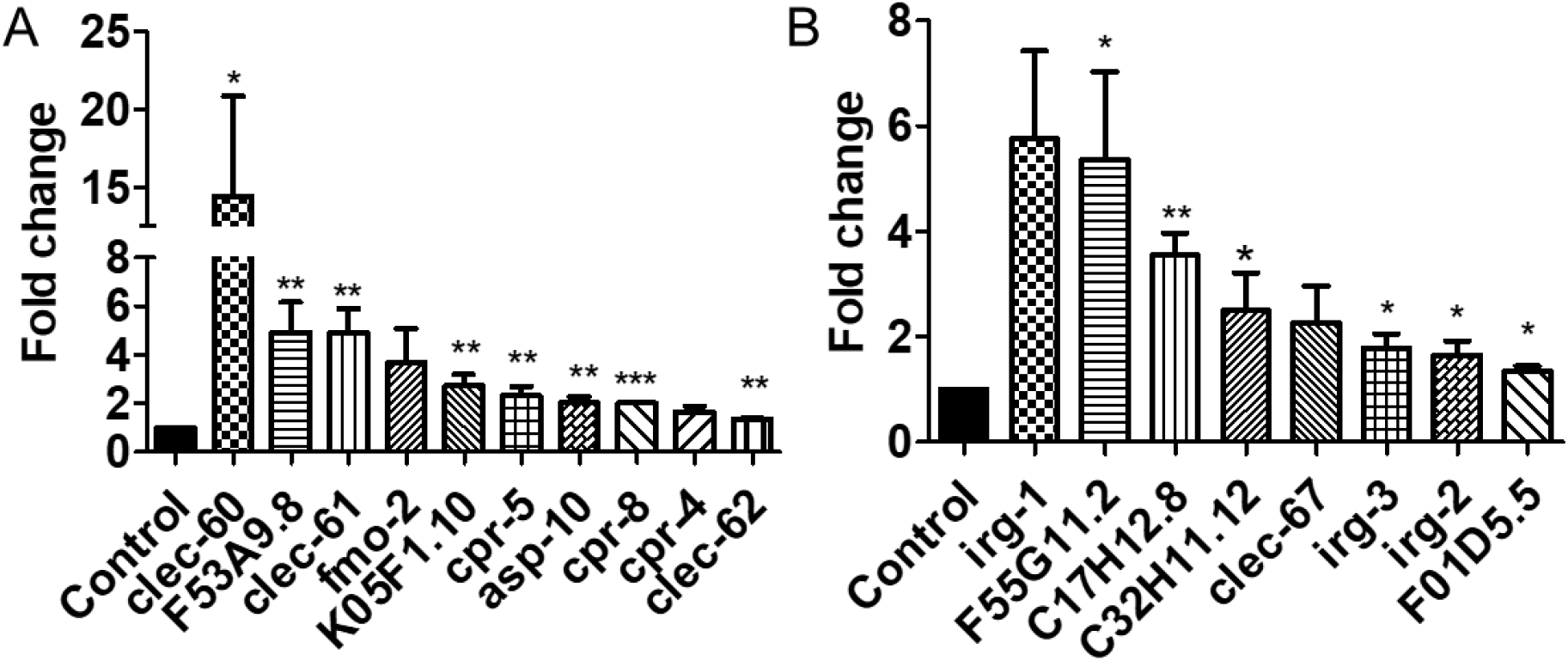
Basal level expression of immune effectors transcripts in *osm-6(p811)* animals compared to wild type. Significance obtained by unpaired *t* test (ns > 0.05, * ≤0.05, **≤0.01, *** ≤0.001).

**Supplementary Figure S4.**
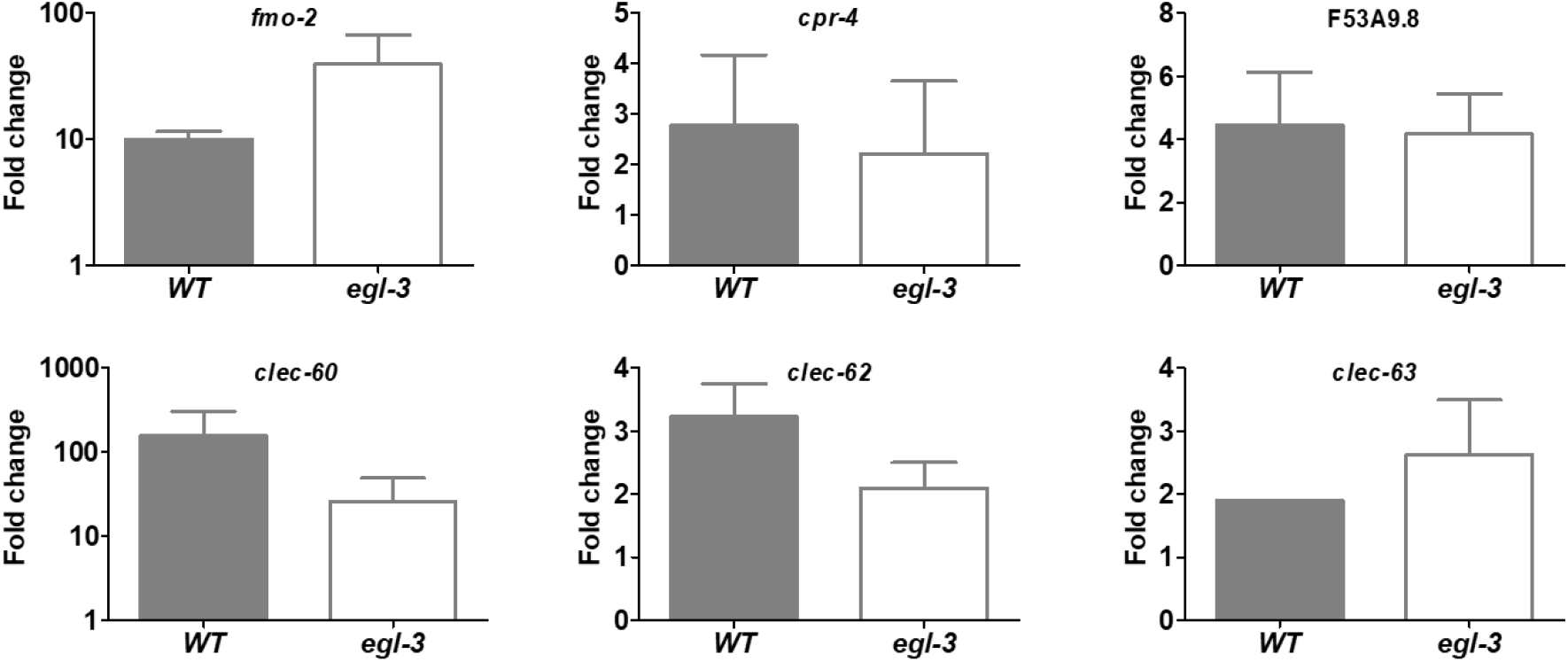
Real time PCR analysis of immune effectors in WT and *egl-3(n150)* animals exposed to *E. faecalis*. (A) *fmo-2*, flavin monooxygenase (B) *cpr-4*, cysteine protease, (C) F53A9.8, hypothetical, (D) *clec-60*, C-Type Lectin, (E) *clec-62*, (F) *clec-63*. Significance obtained by unpaired *t* test (ns > 0.05, * ≤0.05, **≤0.01, *** ≤0.001).

**Supplementary Figure S5.**
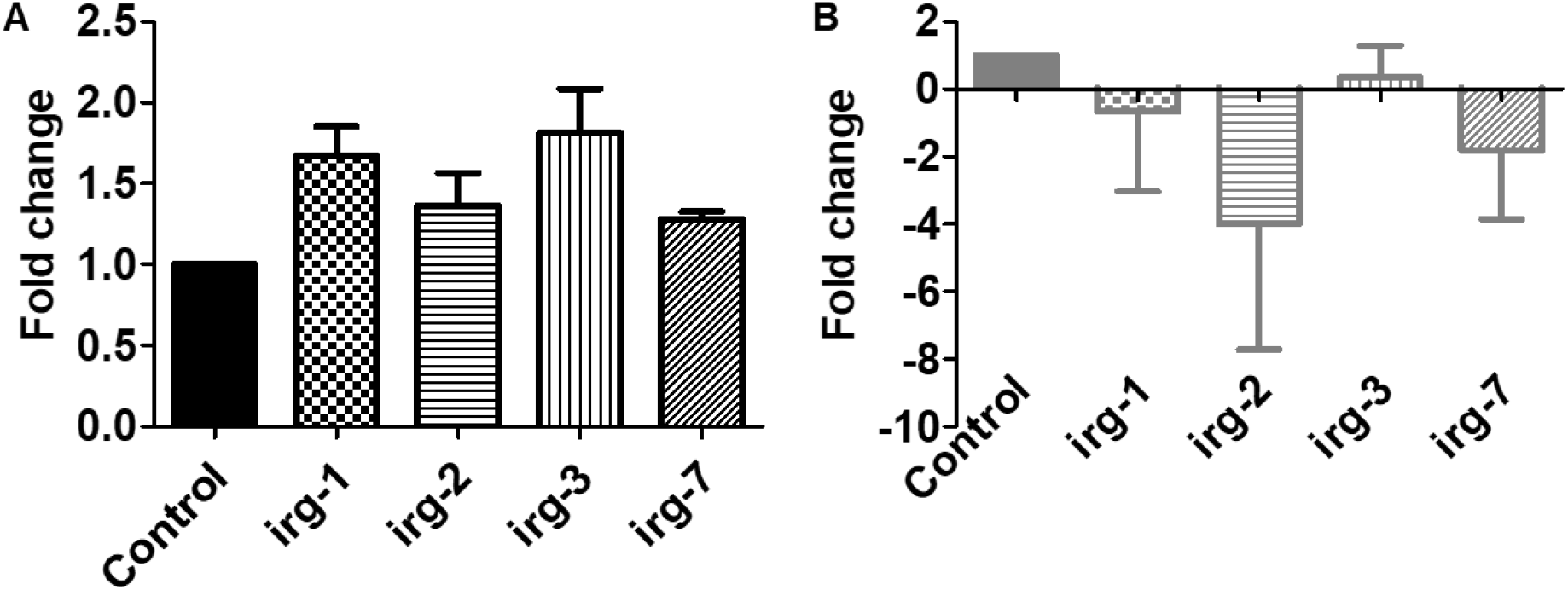
Real time PCR analysis of immune effectors transcripts in WT animals with vector and *fshr-1* RNAi followed by exposure to *P. aeruginosa*. (D) Real time PCR analysis of immune effectors transcripts in *osm-6(p811)* animals with vector and *fshr-1* RNAi followed by exposure to *P. aeruginosa*. Significance obtained by unpaired *t* test (ns > 0.05, * ≤0.05, **≤0.01, *** ≤0.001).

**Supplementary Figure S6.**
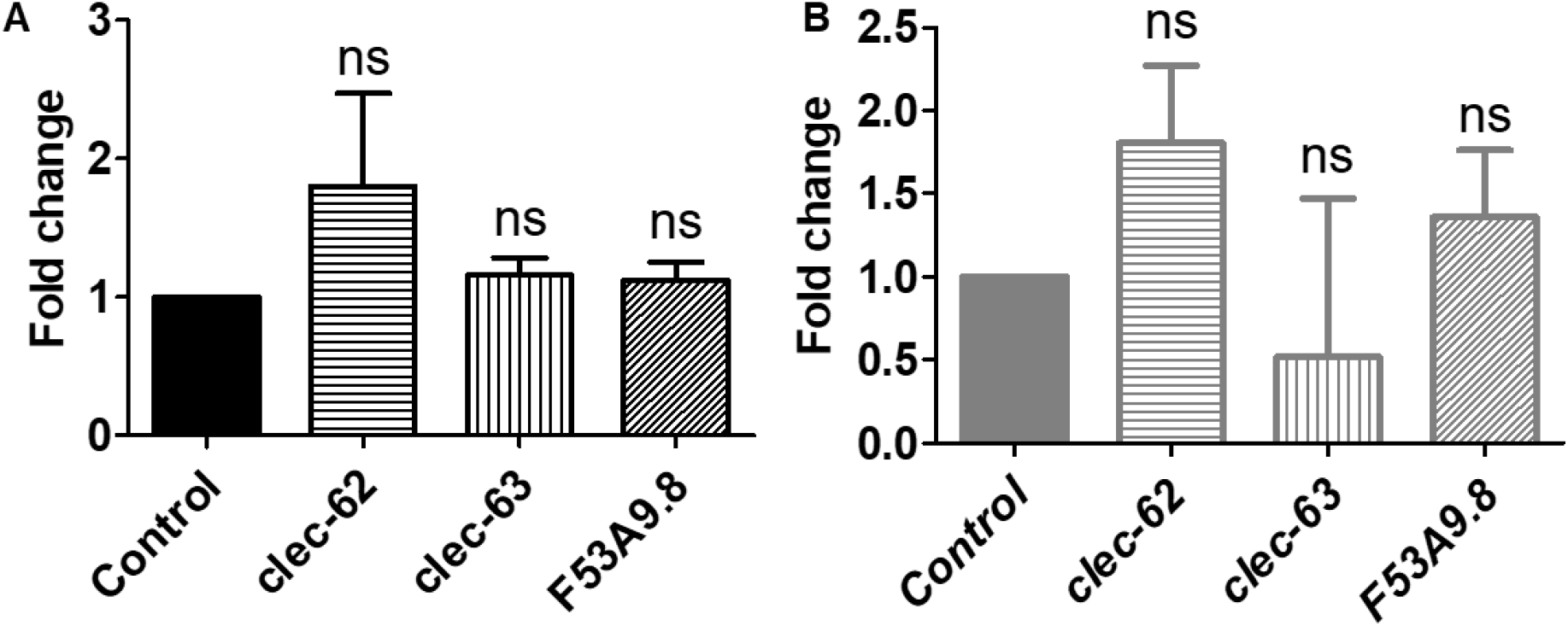
Real time PCR analysis of immune effectors transcripts in WT animals with vector and *fshr-1* RNAi followed by exposure to *E. faecalis*. (D) Real time PCR analysis of immune effectors transcripts in *osm-6(p811)* animals with vector and *fshr-1* RNAi followed by exposure to *E. faecalis*. Significance obtained by unpaired *t* test (ns > 0.05, * ≤0.05, **≤0.01, *** ≤0.001).

**Supplementary Figure S7.**
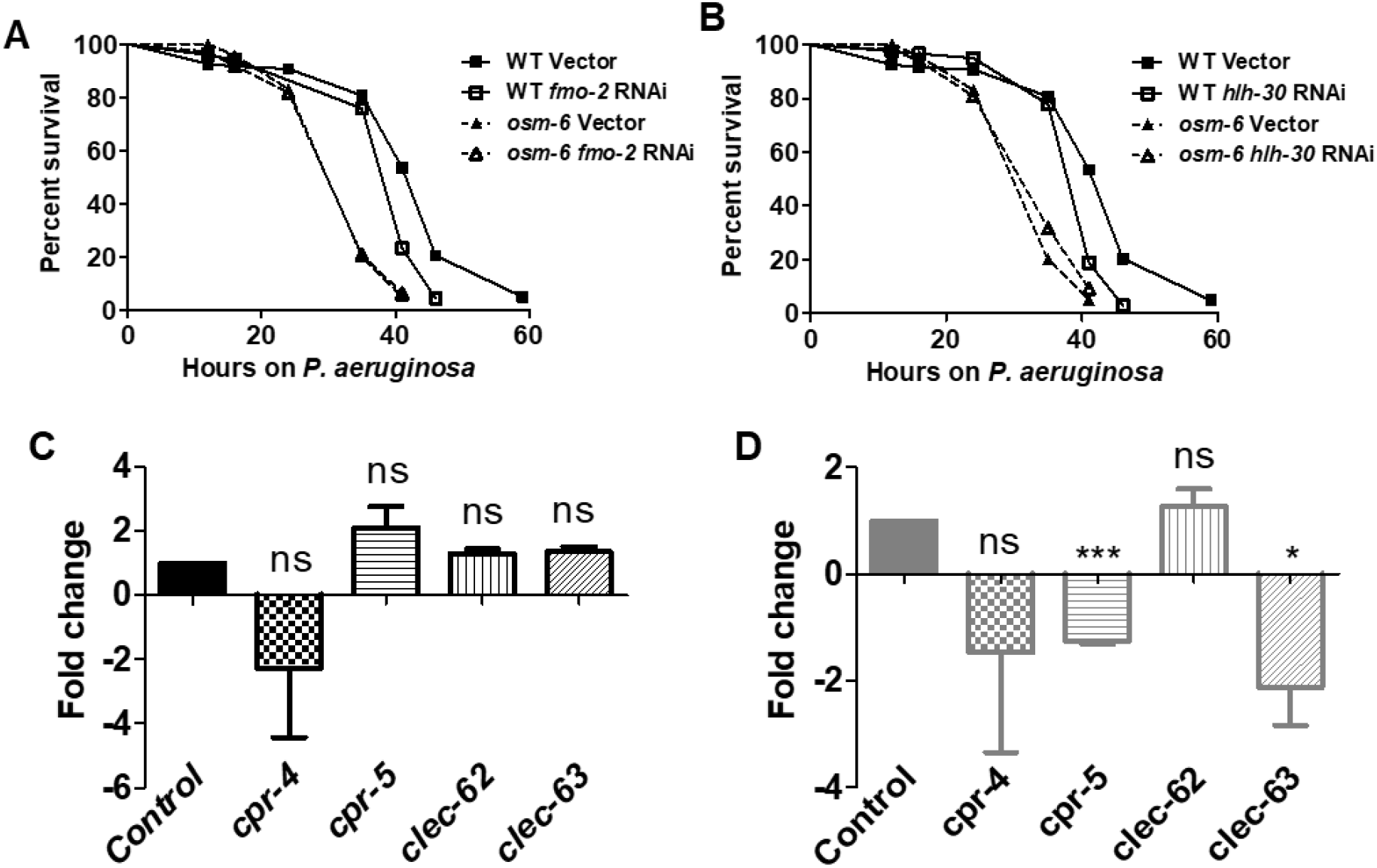
Kaplan Maier survival curve of (A) WT and *osm-6(p811)* animals grown on vector and *fmo-2* RNAi on *P. aeruginosa*. (B) WT and *osm-6(p811)* animals grown on vector and *hlh-30* RNAi on *P. aeruginosa* and scored for survival. (C) Real time PCR analysis of immune effectors transcripts in WT animals with vector and *hlh-30* RNAi followed by exposure to *E. faecalis*. (D) Real time PCR analysis of immune effectors transcripts in *osm-6(p811)* animals with vector and *hlh-30* RNAi followed by exposure to *E. faecalis*. Significance obtained by unpaired *t* test (ns > 0.05, * ≤0.05, **≤0.01, *** ≤0.001).

